# Parametric Deconvolution for Cancer Cells Viscoelasticity Measurements from Quantitative Phase Images

**DOI:** 10.1101/2021.04.06.438595

**Authors:** Tomas Vicar, Jaromir Gumulec, Radim Kolar, Jiri Chmelik, Jiri Navratil, Larisa Chmelikova, Vratislav Cmiel, Ivo Provaznik, Michal Masarik

## Abstract

In this contribution, we focused on optimising a dynamic flow-based shear stress system to achieve a reliable platform for cell shear modulus (stiffness) and viscosity assessment using quantitative phase imaging. The estimation of cell viscoelastic properties is influenced by distortion of the shear stress waveform, which is caused by the properties of the flow system components (i.e., syringe, flow chamber and tubing). We observed that these components have a significant influence on the measured cell viscoelastic characteristics. To suppress this effect, we applied a correction method utilizing parametric deconvolution of the flow system’s optimized impulse response. Achieved results were compared with the direct fitting of the Kelvin-Voigt viscoelastic model and the basic steady-state model. The results showed that our novel parametric deconvolution approach is more robust and provides a more reliable estimation of viscosity with respect to changes in the syringe’s compliance compared to Kelvin-Voigt model.

## I. INTRODUCTION

Examination of a cell mechanical properties becomes an important parameter in cell biology [1]. Typically, the viscoelastic parameters are used to describe particular cells’ mechanical properties using different methods (e.g., atomic force microscopy, optical trapping). Nevertheless, the application of shear stress on adherent cells in a flow chamber setup has become widely used due to the relatively easy setup and promising results (e.g., in [2]). Furthermore, the connection with quantitative phase imaging (QPI) may provide a possibility to evaluate cell deformation after shear stress in a label-free setup [3]. The quantification of shear stress influence on adherent cells is typically based on assessing the cell’s centre of mass as a step response [4], e.g. the reaction of the cell’s movement on the step change of the flow. Due to the hydrodynamic system properties and elastic properties of the tubing and pumping system (typically a syringe), it is impossible to apply a step change of fluid flow (i.e. shear stress) on the cells. There is a noticeable delay in reaching the flow to the set level (in particular in a system with flexible tubing) causing a systematic error in measurements. Minimalisation of this effect may therefore increase the reliability of cell viscoelastic properties estimation. It may make it possible to obtain reproducible results with more compliant plastic syringes (often more easy-to-work with) and significant refinement with rigid ones.

In this paper, we apply the linear system theory to a typical fluid system setup for shear stress application. We apply the deconvolution of the system impulse response to increase the robustness and reliability of the Kelvin-Voigt viscoelastic model. We also compare this approach with direct model fitting and with the estimation of cell shear modulus in steady-state.

## II. METHODS

### A. Cell preparation

Adherent PC-3 prostate cancer cell lines were cultured in Ham’s F12 medium with 7% fetal bovine serum. Themedium was supplemented with antibiotics (penicillin 100 U/ml and streptomycin 0.1 mg/ml). Cells were maintained at 37°C in a humidified (60%) incubator with 5% CO_2_ for 48 hours before exposure to shear stress. RPMI medium with FBS and without antibiotics was used for shear stress induction.

### B. Shear stress system and data acquisition

A simple setup for shear stress application is shown in Fig. 1. A QPI microscope (Telight, Q-Phase) equipped with objective 10x/NA 0.3 was used for acquisition. A programmable syringe pump (Syringe pump Chemyx Fusion 4000) is used to set the flow rate through the tubing connected to the flow chamber (Ibidi µ-Slide VI 0.1 with ibiTreat surface). The actual value is measured by a flow meter (Sensirion SLF3S-1300F) and recomputed to shear stress using the flow chamber’s geometry based on the flow Application Note.

**Fig. 1.**
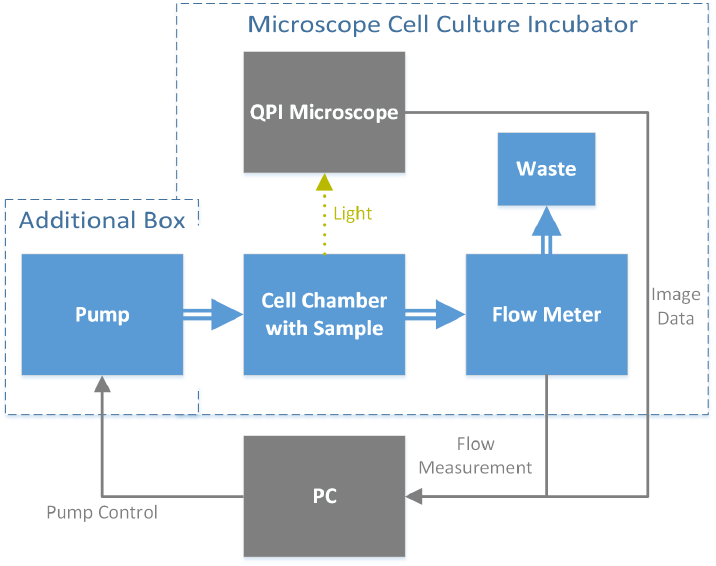
Block scheme of flow system and stress measuring. Blue – flow system generating shear stress in the cell chamber, gray – data acquisition and processing.

### C. Image processing and strain calculation

QPI provides the phase image Φ(*x, y*), which allows to generate a quasi-3D cell projection *d*(*x, y*) via the assumption of a homogeneous refractive index difference Δ*n* (0.02), between the culture media and the cell. Alternatively, under the assumption of homogeneous specific refraction increment, *α* (0.18 *µm*^3^/pg) it allows to generate mass density image *m*(*x, y*) :

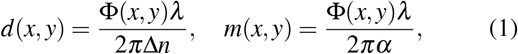

where *λ* is the wavelength of acquisition light. Therefore, both median cell height *b* and cell centre of mass (*CoM*) movement can be measured and used for the calculation of shear strain [4]

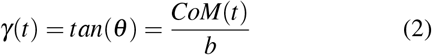

where theta is the shear angle of deformation created by shear stress.

For further analysis, segmentation and tracking of cells in QPI image sequence are required to calculate *h* and *CoM*(*t*) (centre of mass movement in time) for individual cells. If individual cells were close and hard to separate, the whole cluster was used for *b* and *CoM*(*t*) calculation to avoid mass changes caused by segmentation inaccuracies. For segmentation, a stack of video frames was filtered by a 3D median filter to suppress noise in each frame. The segmented cell clusters were obtained by thresholding using a small positive threshold value. Obtained binary images were post-processed by area-based filtering, which removes small objects. Cell clusters touching the border of the actual field of view were removed in each frame (i.e., the whole-cell cluster must be visible). Due to the 3D median filtering and the relatively high acquisition frame rate, the segmented cell cluster forms 3D objects in the video stack image (the cell area in each frame is overlapped with the same cell area in the following frame), which provides tracked cells for following analysis. Afterwards, *CoM*(*t*) and *b* were calculated for each cell cluster.

### D. Strain signal processing and analysis

The cell viscoelastic properties can be modelled by the Kelvin-Voigt (K-V) model, which is described by the parallel connection of the spring (with shear modulus *G*) and damping element (with damping factor *η*). Here we summarize previously applied methods for their estimation, and we describe a novel convolution-based approach for their estimation. The whole analysis process is summarized in Fig. 2. For the K-V model and steady-state model, it is necessary to estimate the cell migration movement from the strain signal and detect the start and end of individual pulses (see Fig. 3a and b with the average pulse signals).

**Fig. 2.**
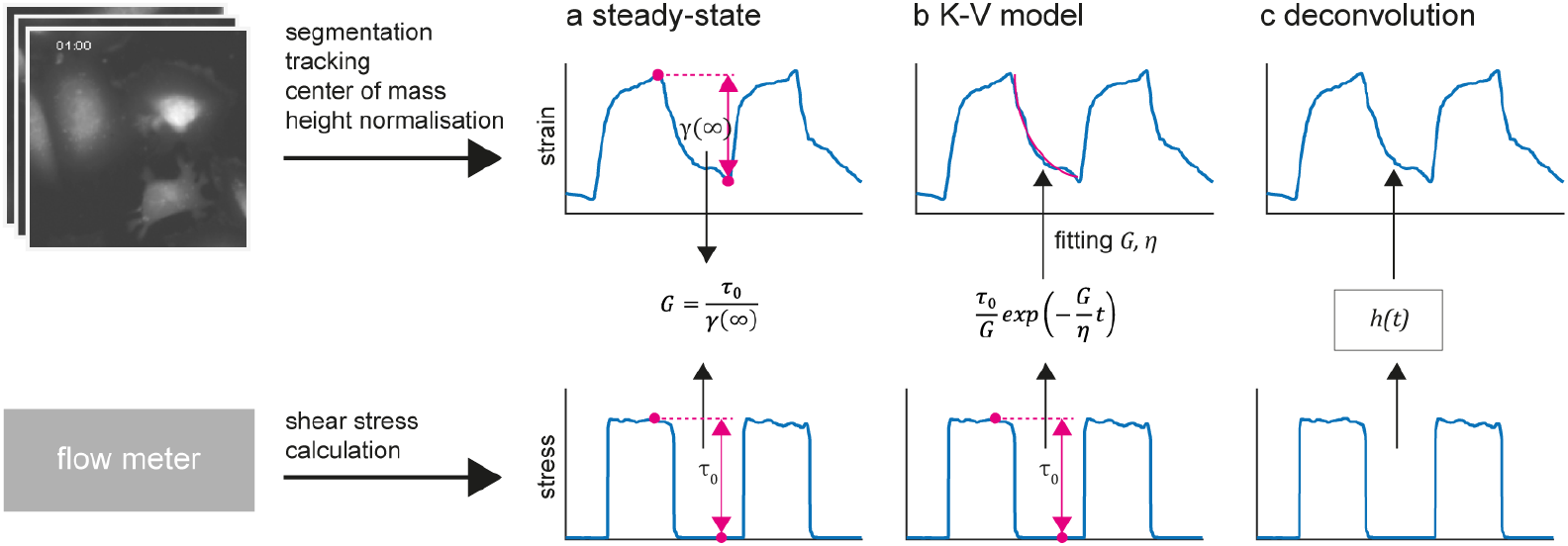
Explanation of the cell viscoelasticity estimation by three different methods. Upper row shows QPI data frames and the resulting shear strain curves over time derived by the used image processing steps (blue signals). Lower row shows calculated shear stress signals corresponding to the measured volumetric flow (blue signals). The basic principles of steady-state method (a), Kelvin-Voigt model fitting (b), and parametric deconvolution method (c) are shown. Pink colour and black arrows show the needed measurement and resulting parameters/signals.

**Fig. 3.**
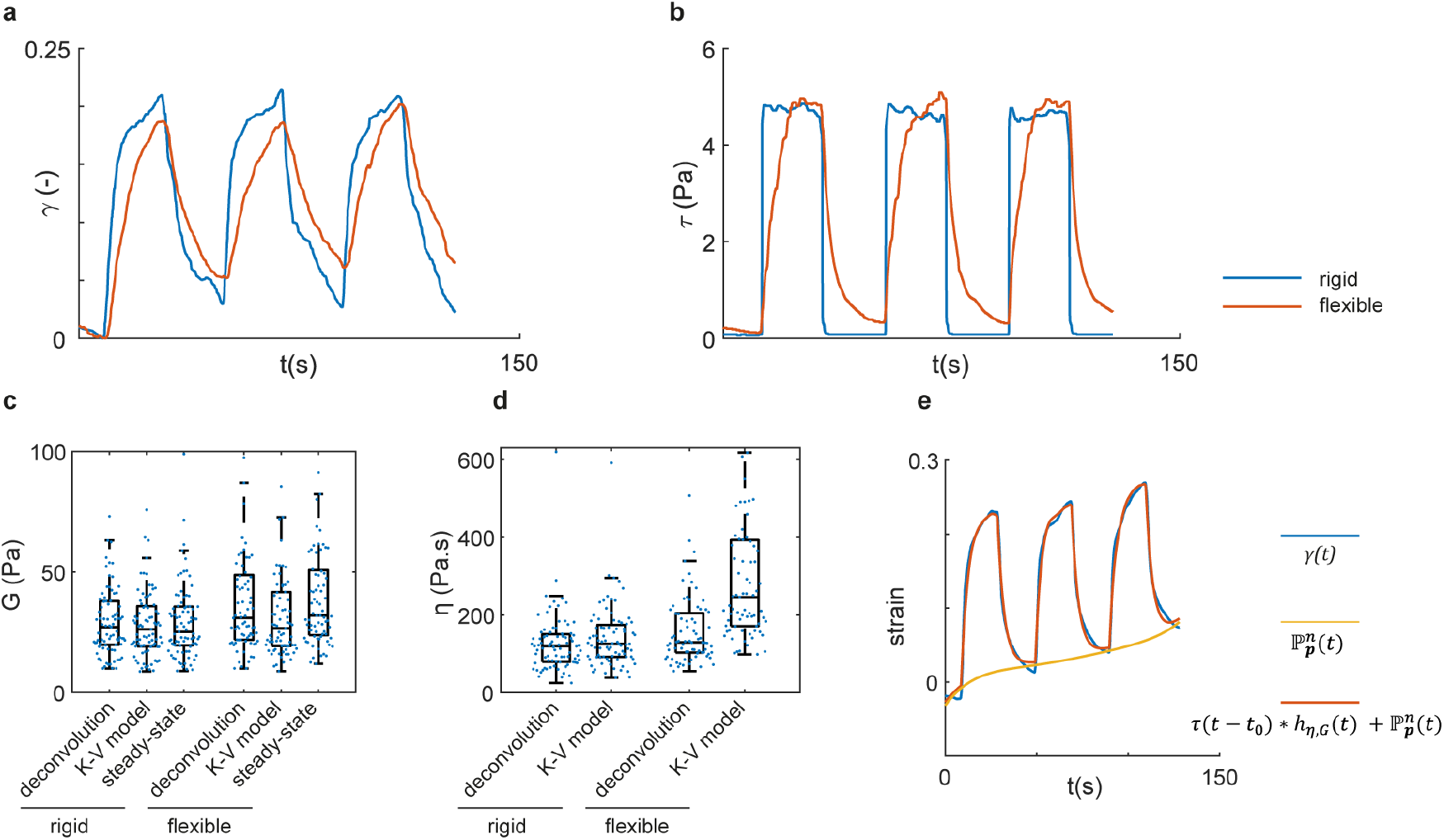
Achieved results and comparison of the influence of different syringe use and different estimation methods. The shear strain signals obtained as an average form all measured cells (a) are shown. The blue signal comes from the ‘rigid’ glass syringe and red signal comes from ‘flexible’ plastic syringe. Similarly, the average shear stress signals are shown in (b). The comparison of achieved estimated values of cell shear modulus (c) and cell viscosity (d) using different flow systems and different models is shown. Blue dots – estimated values of different cells/cell clusters, box edges – 75th/25th percentil, middle line – median value, whiskers – maximal/minimal value from 1.5 times of the inter-quartile range. Example of the shear strain signal fitting using the parametric deconvolution model (e). Blue signal – measured shear strain signal, yellow signal – fitted polynomial model of centre of mass changes caused by cell migration, red signal – estimated shear strain signal combining convolutional distortion modelling and cell migration modelling.

The migration movement still occurred in the direction of the applied flow, which biased the CoM values. To suppress this effect, the movement signal was removed by 5-th order polynomial fitting and signal subtraction. The sample differences were calculated, and the peak detector was applied (with a selected threshold and minimum peak distance). The sample differences of the shear stress signal were calculated, and the peak detector was applied to detect the start and end of the pulse. Because of the delay between the generated shear stress signal and the CoM response, the detected pulse start and end positions did not exactly correspond to the extrema in the CoM signal. The correction of the position was done by a locally restricted maxima detector. Positions of the start and end of the pulses were then used for the calculation of differences in the steady-state model and for the selection of a part of the signal for K-V model fitting.

For the cell viscoelasticity estimation, the three following methods/models have been used:

#### 1) Kelvin-Voigt model

The response (strain) of the cell to the unit step function of the shear stress can be described by a decaying exponential model:

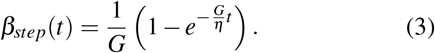

The applied shear stress has typically specific value(s) *τ*_0_ and therefore the response can be modified as:

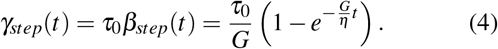

The left-hand side of this equation corresponds to the shear strain (eq. 2). The values of *τ*_0_ are known and set during the experiments; thus, the model parameters (cell shear modulus *G* and viscosity *η*) can be obtained by simple curve fitting. The model was fitted by a robust bisquare method [5] optimized by Levenberg–Marquardt algorithm (nonlinear least-squares) [6]. Additionally, we have included the fitting of the parameter *c*, which is added to the fitted exponential function and serves for the cell movement correction because the previous cell movement correction is not precise enough.

#### 2) Steady-state model

The K-V model for steady-state condition provides a simple way to estimate the shear modulus *G*. If we let *t* → ∞, then the expression in the bracket of eq. 4 is equal to 1 and *G* can be estimated as:

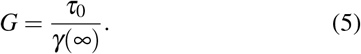

Thus, the determination of *CoM* shift in steady-state can provide a simple shear modulus estimation. A pulse length of 20 seconds was used in our experiments, sufficient for achieving steady-state with the required precision.

#### 3) Parametric deconvolution

The single cell can be considered as a linear system, which can be described by an impulse response *h*(*t*) computed as a differentiation of the step response (eq. 4):

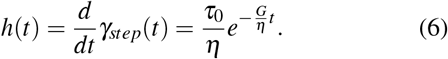

This impulse response can be further used to model the origin of the cell’s shear strain as *γ*(*t*) = *h*(*t*) ∗ *τ*(*t*). This general expression for the arbitrary shear stress waveform *τ*(*t*) might be applied in our model identification problem.

Let us formulate an optimization task with the following cost function:

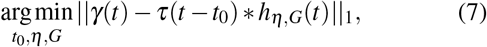

where *h*_*η,G*_(*t*) is a parametric impulse response (eq. 6) with unknown parameters *η* (cell viscosity) and *G* (cell shear modulus), *τ*(*t* − *t*_0_) is a shear stress applied on the cell (generally shifted by unknown shift *t*_0_) and *γ*(*t*) is the measured cell displacement normalized by its height (i.e., shear strain). Nevertheless, we observed that low-frequency ‘trend’ signal of cell migration movement is typically present in the measured data (see Fig. 3e) and this additional signal distorts the optimization process. To cope with this problem, we extended our optimization problem by additional terms composed of the polynomial 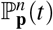 fitting:

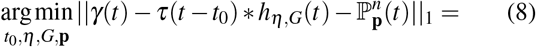

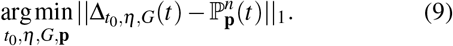

During the optimization, the polynomial fit (with preset polynomial order *n*) of the actual remaining signal 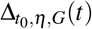 is performed and the parameter vector **p** of the polynomial fitting curve is found.

## III. RESULTS AND DISCUSSION

In our experiments, we tested the influence of the syringe on the estimated cell parameters. We observed that different syringes significantly influence the waveform of the flow, particularly during the fast flow changes, due to the different syringe compliances. The first syringe was FORTUNA Optima glass syringe 20 ml (referred as ‘rigid’), and the second one was Braun Original-Perfusor plastic syringe 20 ml (referred as ‘flexible’). The ‘flexible’ syringe has the rubber seal of the plunger, which increases its compliance. Stimulation in our experiments consists of 3 subsequent square pulses, each lasting lasting 20 seconds, with a 20 seconds gap in between. The magnitude of the pulses was set to achieve shear stress equal to 5 Pa, which is in the physiological range (i.e., 0 Pa – 10 Pa, according to Malek et al. [7]). This preset shear stress value is sufficient for cell deformation, which causes the movement of the CoM position of the particular cells. To evaluate the methods of strain signal analysis from Section II-D, i.e., K-V model, steady-state model, and parametric deconvolution model, we have measured four fields of view with both ‘rigid’ and ‘flexible’ syringes with the proposed setup (Section II-B). The number of cells was 75 and 83 for ‘flexible’ and ‘rigid’ syringe, respectively. Average of the shear strain and shear stress signals of all cells for both syringes is shown in Fig. 3 a and b, respectively. It can be seen that the shear stress signal from ‘flexible’ syringe system has considerable difference compared to the optimal step function and thus, without any compensation of this effect, the estimated cell parameters might be significantly distorted.

The K-V model method using exponential fitting is based on the assumption that the shear stress stimulation is a step function. This assumption approximately holds for the ‘rigid’ syringe; however, as we can see before, for the ‘flexible’ syringe, this assumption does not hold, and it will lead to the falsely increased value of the estimated viscosity *η*. This situation might also occur if there is an air bubble inside the syringe or tubing system or if the tubing itself is not rigid enough.

The steady-state model considers only the steady values, and therefore, if the time to achieve the steady-state is long enough, it will not be influenced by the elasticity of the flow system. Consequently, this model cannot be applied in the case of fast stress changes together with a too elastic flow system. Another disadvantage of this model is its impossibility to estimate the viscosity parameter *η*.

Finally, the proposed parametric deconvolution method works for various input shear stress signals. It predicts the similar viscosity *η* for both rigid and flexible syringes due to eliminating the influence of the syringe and tubing system (see Fig. 3d).

As expected, the estimated shear modulus *G* is similar for ‘rigid’ and ‘flexible’ syringes for all methods (as can be seen in Fig. 3c). For viscosity estimation, however, the K-V model leads to highly distorted results – with a median viscosity of 125.8 *Pa* · *s* for a rigid and 246.0 *Pa* · *s* for the flexible syringe (an increase of 95%). However, the viscosity for the deconvolution method incenses only slightly from 119.8 *Pa* · *s* to 129.2 *Pa* · *s* (8% increase). Results for both methods are shown in Fig. 3d.

This result shows that the basic linear model for the syringe and tubing system can efficiently remove their influence via parametric deconvolution. Furthermore, this approach is not based on the necessity of the input step function, but it might be applied for arbitrary shear stress waveforms. Another advantage is the novel approach for the elimination of the migration movement, which causes low-frequency distortion of the measured CoM(t) signal. We showed that the polynomial fitting of the remaining signal 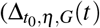, from eq. 9) can efficiently model and remove influence of migration movement (see a yellow curve in Fig. 3e).

## IV. CONCLUSIONS

We have compared three different methods for the estimation of viscoelasticity parameters using QPI microscope and flow-based shear stress application. The proposed parametric deconvolution method, unlike the other two methods, is able to robustly estimate both the shear modulus and viscosity. Its robustness was tested on a setup with rigid and flexible syringes, which highly influences the tubing system response, where the estimated viscoelasticity was similar for both setups. The whole setup is able to acquire the image and flow data. The proposed image processing pipeline is able to extract shear strain for individual cells, which can be used for the estimation of cell viscoelasticity parameters.

## ACKNOWLEDGMENT

This work is supported by the Czech Science Foundation project no. 18-24089S.

## Notes

### Competing Interest Statement

The authors have declared no competing interest.

